# From Functional Organization to Vascular Structure in the Primate Visual Cortex Using Ultrasound Imaging

**DOI:** 10.64898/2026.08.18.745404

**Authors:** Erwan Dessailly, Ignacio Alcala, Matthieu Provansal, Rafik Arab, Nicolas Zucker, Florian Fallegger, Pierre Pouget, Mickael Tanter, Serge Picaud, Fabrice Arcizet

**Author notes:** These authors contributed equally (Erwan Dessailly, Ignacio Alcala, Matthieu Provansal). These authors contributed equally (Serge Picaud, Fabrice Arcizet).

## Abstract

The primary visual cortex (V1) provides an exceptional structure to assess information encoding in the cortex thanks to its highly controlled visual input. However, optical measures and electrophysiology are limited to very superficial layers while fMRI provides low spatial resolution. Taking advantages of functional ultrasound imaging (fUSi), we examined large volumes of V1 in two awake non-human primates. We showed the conservation of retinotopic maps within deep sulcal regions and across cortical layers. We then assessed the cortical point image and compared it to its vasculature distribution through ultrasound localization microscopy (ULM). We further identified the smallest vascular compartment underlying the signal. Together, these results establish fUSi, complemented by ULM, as a minimally invasive mesoscale approach for resolving functional organization of the cortex in deep sulci.

**Teaser:** Functional Ultrasound imaging (fUSi) on awake nonhuman primates reveals retinotopic functional maps to point image and high-resolution images of the vasculature system.

## Introduction

Sensory and motor information are processed through a network of multiple interconnected cortical areas arranged hierarchically based on laminar connectivity patterns (*1*, *2*). These cortical structures are composed of 6 layers interconnected by pyramidal neurons extending vertically throughout the different layers. This apparent vertical processing appears organized around lobular microvascular units forming a motifs embedded within the cortical mosaic (*3*). This anatomical vascular unit is composed of arteriole and venule with reported ratios ranging from ∼2.6:1 (∼12 arterioles/mm² and ∼5 venules/mm²) in histological studies (*3*) to ∼1.4:1 (∼4.1 arterioles/mm² and ∼3.2 venules/mm²) in a recent in vivo 7T MRI study (*4*). Indeed, this anatomical vascular unit has not yet been clearly imaged in vivo nor linked to a functional unit.

To investigate the structural and functional organization of the cortex, we have taken advantage of the primate primary visual cortex (V1) which can be finely stimulated visually. Indeed, V1 exhibits a retinotopic organization, where each stimulus position in visual space aligns with a matching site first on the retina to the cortex. Retinotopic mapping can be achieved across multiple spatial scales using complementary imaging techniques. fMRI provides whole-brain coverage at millimeter resolution (*5*, *6*) revealing on non-human primate mesoscopic features such as ocular-dominance orientation domains (*7–10*). Optical methods (intrinsic optical signal as IOS and two-photon) offer much higher, near-cellular resolution but are limited to superficial cortical layers (< 700 µm, (*11–14*). Ultra-high field MRI (≥ 7T) has overcome this gap by enabling non- invasive mesoscopic mapping across the entire cortex and across cortical layers, including deep sulci, with improved spatial specificity using spin-echo sequences (*15*, *16*).

Beyond retinotopy, visual cortex organization is also characterized by the point image defined as the cortical space activated by a single point in visual space (*17*). Basically, one can easily computed it as the product of the cortical magnification factor (CMF in mm/deg) and the population receptive fields size (RFs, in degrees). The CMF describes how visual space is non-uniformly represented, with the fovea occupying a larger cortical area than the periphery whereas the RF reflects the spatial sensitivity of individual neurons (*18*). As the eccentricity increases the neuronal RF becomes larger and the CMF decreases, pointing toward a constant point image of 2 to 4 mm (*17*). It is intriguing to consider how this point image relates to the vascular network unit described above.

Functional ultrasound imaging (fUSi) has recently emerged as a novel imaging modality combining deep-tissue penetration with high sensitivity to measure the neurovascular coupling. fUSi offers high-resolution (∼100 µm), deep-tissue functional mapping by measuring cerebral blood volume changes, enabling retinotopic and activity maps in primates (*19–21*). When compared to fMRI, fUSi is a highly portable and easily transportable technology but it requires a cranial window. The same machine can also generate Ultrasound localization microscopy (ULM) to reveal ultra-high-resolution vascular 3D maps (∼27–60 µm), (*22–24*). Here, thanks to this novel fUSi technology, we question the conservation of the retinotopic organization across different cortical layers (infragranular versus supragranular) on primate V1. Furthermore, combining fUSi and ULM on the same spot enables us to address the relationship between the cortical point image and the smallest functional vascular unit.

## Results

In this study, we used a linear ultrasound probe providing functional activation maps with a spatial resolution of 100 × 100 × 400 µm, imaging a plane spanning 14 mm in width and up to 20 mm in depth. The recording chambers placed above the visual cortex of two non-human primates were located on adjacent yet distinct cortical regions, ensuring that the two fUSi sagittal imaging planes were contiguous along the visual cortex (Figure 1A). For Monkey L, its chamber was positioned over the right lunate sulcus (ML: +7 mm; AP: -10 mm), and for Monkey E over the right calcarine sulcus (ML: +7 mm; AP: -22 mm). Both recording chambers enable us to image a large portion of the V1 area along the calcarine sulcus, including both superficial and deep layers in Monkey E, and to capture the superficial layer of V2 and the deep layers of V2 and V3 in Monkey L (Fig. 1A). To correlate the variations of CBV and the presentation of visual stimuli, we used the General Linear Model (GLM) approach, as commonly applied in fMRI (*25–27*). We first characterized the hemodynamic response function (HRF) for each accessible region of the visual cortex in both monkeys (E and L; Fig. 1B). For each HRF, we averaged the cerebral blood volume responses evoked by full-field checkerboard stimulation (see Methods) and fit the resulting time course with an inverse-gamma function. For Monkey L, we reused the HRF from a previous study (*20*), while for Monkey E, we applied the same protocol to derive an animal and region-specific HRF (Fig. 1B). Using these HRFs, we generated predicted CBV signals by convolving the stimulus presentation timestamps with the HRF. These predictions were correlated, voxel by voxel, with the fUSi time series to generate z-score activation maps. In our pipeline, the inputs are the Doppler image time series and the stimulation timestamps; the stimulus sequence is convolved with the HRF to yield a CBV prediction, which is compared to each voxel’s signal and converted to z-scores. The resulting z-score maps were then used to derive functional maps (Fig. 1B, right panel).

**Figure 1.**
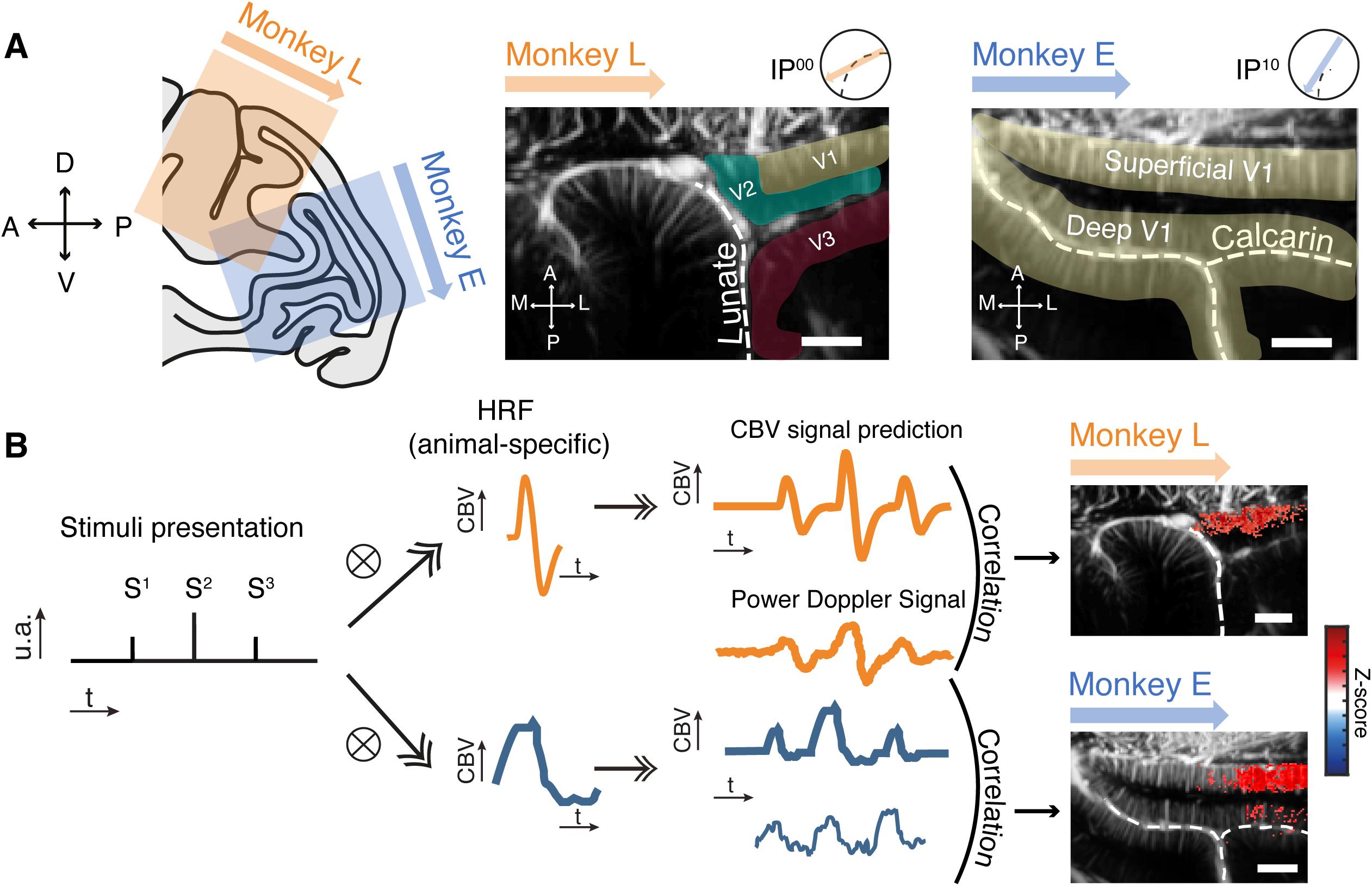
Experimental set-up and General linear model used for fUSi analysis. **(A)** Anatomical positions of the recording chambers for Monkey E and Monkey L. The left panel represents a longitudinal view of a rhesus brain at ML +7 mm derived from an atlas; we also indicated the field of view accessible through the recording chamber for each animal (in orange for Monkey L centered on the lunate sulcus, and in blue for Monkey E centered on the calcarine sulcus). The right panels represent the anatomical images obtained with the ultrasound probe. The dashed lines indicate the lunate sulcus for Monkey L, and the calcarine sulcus for Monkey E. Scale bars are 2 mm. The colored arrows symbolize the position of the ultrasound probe inside the recording chamber (the black circle), and the position of the arrowhead (on the top of the anatomical image) indicates the orientation of the image. **(B)** GLM-based generation of functional maps. The prediction was convolved for functional ultrasound imaging data with an animal-dependent hemodynamic response function. We first determined the HRF for each animal. We then correlated the CBV signal prediction to raw power doppler signal. We represented all the significant Z-score voxels superposed to the corresponding anatomical image.

### Retinotopic mapping of primate V1 in multiple imaging planes

To investigate the retinotopic representation of the primary visual cortex, we decomposed our visual field in eccentricity and polar angle. All stimuli were presented in the left visual hemifield while the animal maintained central fixation within a 2.5° window around the fixation point, allowing us to separately map polar angle (angular position around central fixation) and eccentricity (distance from fixation) selectivity.

To study the different polar angle’s locations, we divided the left visual hemifield into a set of 12 pie-slice visual stimuli; each stimulus covered each subtending 15° of polar angle from 2 to 15 degrees of visual angle (DVA) of eccentricity, (Figure 2). In other words, the 12 wedges (12 × 15° = 180°) tiled the entire left hemifield and restricting the wedges to 2–15° DVA excluded the foveal region while keeping eccentricity coverage comparable across polar-angle conditions. We investigated the retinotopic organization of the primary visual cortex by imaging 12 equiangular imaging planes centered on the circular recording chamber (see Figure S1, one imaging plane “IP” every 30 deg), providing broad spatial sampling of the visual cortex. The retinotopic selectivity for polar angles is visible in Figure 2A as the resulting activation patterns shown on Monkey E for six of the twelve polar angles varied systematically with stimulus location. For instance, stimuli presented in the upper left visual quadrant evoked maximal responses in the most ventral portion of the superficial layers of V1 (image #2), while stimuli in the lower quadrant activated more dorsal regions (image #5), indicating a clear spatial mapping of polar angle preference. This dorsal–ventral shift is consistent with the expected inversion of the visual field representation in V1 (upper visual field represented ventrally; lower visual field represented dorsally).

**Figure 2.**
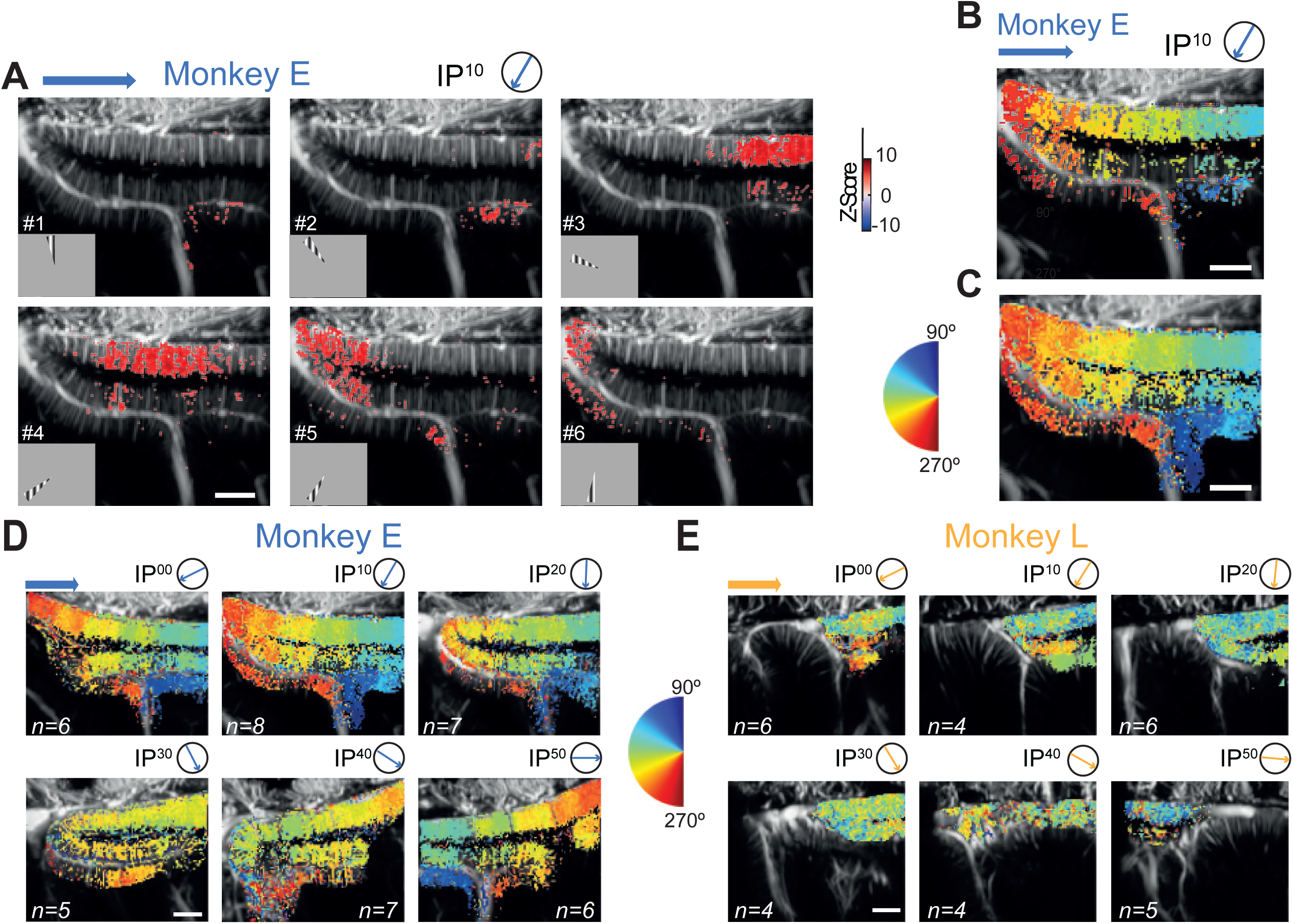
Retinotopic maps based on different polar angles. **(A)** Activation maps according to t6 different polar positions of the visual stimulus (here, a portion of a circular vertical grating). The circle schematized the recording chamber and the blue arrow the position of the ultrasound probe. The head of the arrow indicates the orientation of the imaging plan (here IP10). Each image represents the activation map for a particular polar angle (90 to 240, each 30°). The color code indicates the value of the z-score. **(B)** A single session polar angle map for one imaging plane (IP10). Polar angle map reconstructed from the activations in A, obtained in a single session for 12 different polar angle stimulations (∼20 trials per polar angle). **(C)** Mean polar angle map for one imaging plane (IP10). We averaged the total number of sessions (n=8) voxel by voxel to obtain a retinotopic map for one specific imaging plane after spatially aligning all individual retinotopic maps per session. **(D and E)** Mean polar angle maps for other imaging planes for Monkey E **(D)** and Monkey L **(E)**. Polar angle maps for polar positions for 6 different imaging planes (IP00 to IP50). For each imaging plan (schematized by the circle and the arrow), several imaging sessions were pooled together. Scale bars: 2 mm.

To summarize activity across all stimuli within a session, we generated composite maps to reveal the phase-encoded retinotopy in which each voxel was assigned the color corresponding to the stimulus that elicited the highest Z-score (Fig. 2B). Thus, these “winner-takes-all” maps directly indicate, for each voxel, the polar-angle condition producing the strongest response. Comparable polar-angle preference maps have also been reported with fMRI in non-human primates using phase-encoded wedge/ring paradigms, supporting the interpretation of these maps as retinotopic selectivity rather than stimulus-specific idiosyncrasies (*28*, *29*). We repeated several times (n= 3 to 7 sessions) and these session maps were then spatially aligned based on the anatomy and averaged across all independent recording sessions per imaging plane (see Methods), resulting in mean polar angle maps (Fig. 2C). Averaging across sessions improved map stability and reduced session-to-session variability, yielding a more robust estimate of polar-angle preference within each plane. (Fig. S3).

We then tested V1 representation of eccentricity using an approach analogous to that used for polar angle mapping (see Figure S2). We presented a series of 2 DVA thick six hemi concentric arc bands centered on the central fixation point spanning eccentricities from 2 DVA to 14 DVA in the left visual field. Using fUSi, we obtained clear, stable and distinct retinotopic maps across different regions of primary visual cortex along anatomical sulci such as lunate and calcarine (see Figure S3 et S4). We observed an eccentricity distribution aligned with these anatomical sulci, with central vision (low eccentricities) being represented in the superficial fold for both animals (Monkey E and Monkey L), and toward peripheral vision in deeper fold (Fig. S2).

### Layer specification of retinotopic maps

We next aimed to determine whether functional ultrasound imaging could resolve the mesoscopic functional architecture across the cortical thickness. To investigate how retinotopic preferences vary across cortical layers and along the medio-lateral axis, we performed a laminar analysis of the two most superficial cortical folds for all imaging planes in both monkeys. Each fold was subdivided into three laminar ensembles: supragranular layer (L1–L3), granular layer (L4), and infragranular layer (L5–L6), as defined in the Methods. For each layer group, fUSi signals were averaged along the z-axis, and the resulting maps were projected onto the cortical surface across successive imaging planes (Fig. 3A). Data from adjacent planes were then interpolated at the in-plane resolution of 400 µm to generate continuous top-view projections of retinotopic selectivity (Fig. 3B and 3C, for eccentricities and polar angle respectively). For Monkey E, all laminar projections revealed a consistent medio-lateral increase in preferred eccentricity and a clear antero-posterior gradient in polar angle preference, confirming the previously observed spatial organization. In contrast, Monkey L displayed no apparent spatial segregation of eccentricity or polar angle in any laminar ensemble around the lunate sulcus. To further assess intra-laminar variations, we compared eccentricity and polar angle preferences across depth for voxels sharing identical (x,y) coordinates and significant selectivity in all three laminar projections. The distributions of these differences are shown adjacent to the chamber representations. Due to high number of voxels, we noticed a slight but statistically significant increase in preferred eccentricity with cortical depth in V1 for both monkeys with similar mean eccentricities per layer. For Monkey E, mean averaged eccentricities varied from 5.2 to 5.5 DVA for superficial V1 and 9.5 to 10.2 for deep V1 although mean averaged eccentricities varied from 6.1 to 6.6 DVA for V1 and 6.0 to 6.4 for V2 for Monkey L (see Supplementary Table 1 and Supplementary Table 2 for descriptive statistics). Concerning polar angle selectivity, the layer 4 displayed a modest shift in preference toward the lower visual field compared to supragranular and infragranular layers superior V1 fold on Monkey E and for V1 area on Monkey L. However, this pattern was not observed in the inferior V1 fold or in V2, suggesting that laminar modulation of polar angle preference may be region-specific or subject to individual variability. Given that both the eccentricity and polar angle intra-laminar variations we report are subtle (below 1 DVA, see Supplementary Tables 1 and 2) and that receptive fields are between 1 to 5 DVA in diameter (*5*), we interpret that the retinotopic preference was conserved across cortical layers. Altogether, these findings indicated that V1 retinotopic architecture is stable across the cortical laminae and varies only in an orthogonal direction of the vessels. We then confirmed that fUSi effectively captures the columnar organization of V1, providing a mesoscopic resolution comparable to that achieved with ultra-high field MRI.

**Figure 3.**
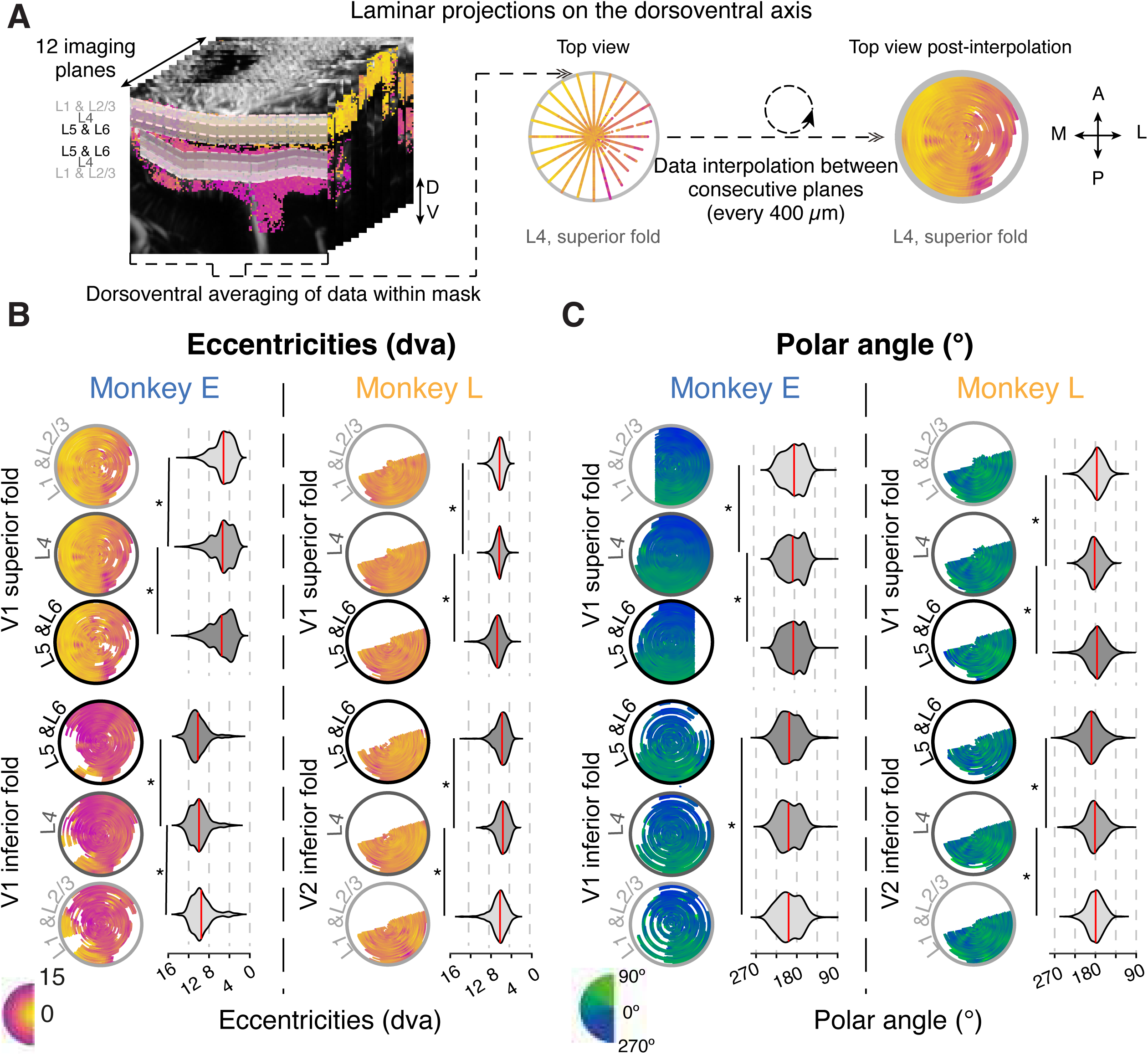
Laminar retinotopic maps. **(A)** Methodology. All retinotopic maps are averaged on the dorsolateral axis in three laminar ensembles (supragranular: L1 & L2/3; granular : L4 ; infragranular : L5 & L6) defined in all cortical folds. For each laminar ensemble, data are interpolated between successive imaging planes at the fUSi spatial resolution (400 µm). **(B)** Eccentricity representation through a cortical layer. Left column: top view of medio-lateral maps of preferred eccentricities for all laminar ensembles in each cortical fold (supragranular, granular and infragranular). Right column: distribution of eccentricities for common voxels between all laminar ensembles within a cortical fold. The two left columns represent retinotopic map for Monkey E although the two right ones for Monkey L. **(C)** Same representations but for preferred polar angles. (*) indicate a p-value <0.001 one-sided paired t-test with Benjamini correction.

### Study of point image and cortical magnification factor in V1

We then investigated with fUSi how the size of a stimulus affects the spread of cortical activation allowing an estimation of the point image and cortical magnification factor (CMF) in one imaging plane. The CMF was obtained by presenting sections of annulus of 6 different sizes (Fig. 4A) while we imaged a single imaging plane (IP^10^ and IP^00^ for Monkey E and Monkey L respectively). For both animals, stimuli were presented at two eccentricities (6° and 11° of DVA) and at two angular positions (191° and 213°), as these locations yielded well-centered activations within the imaging plane. For each session, we generated activation maps for every size of stimulus as represented in Figure 4A (stimulus centered at 11 DVA of eccentricity and at a 213°angle). As expected, activated areas increased with stimulus size. Moreover, by overlaying these activation maps, we noticed that activation maps were confined to the same cortical layer location for Monkey E spreading along the medio-lateral axis whereas in Monkey L they spread across superficial and deep layers of primary visual cortex (Figure 4B). We then plotted the spread of activity as a function of the square root of stimulus area, both for single sessions per position (see Figure S5) and averaged per eccentricity (Fig. 4C). Finally, we computed the CMF for each eccentricity by fitting a linear regression for both animals (see methods). In both monkeys, we observed a linear relationship between stimulus size and cortical activation (Fig. 4C). Consistently for both monkeys, stimuli at an eccentricity of 6 DVA elicited a wider spatial spread of cortical activity than those at an eccentricity of 11 DVA—approximately 1.5× more activity for a 13.57 deg² stimulus (highest size stimulus at eccentricity 6) in Monkey E—consistent with the cone and ganglion-cell densities decreasing according to a fovea to periphery gradient (*30*). Overall, for higher size of stimuli the fUSi measurements were reliable with more variability for Monkey L. For instance, activity resulting from small stimuli (0.6 and 1.5 deg²) were already spreading in width along the upper fold for ∼1 to 2 mm (depending on eccentricity, Fig. 4B). Already for the second size of stimuli, the upper fold of Monkey L was almost fully activated. This correlates with the standard deviation of the retinotopic maps (see Figure S4) and point to a non-selectivity of this upper fold. Moreover, for stimuli bigger than 6-7°² much of the observable cortex was recruited, producing a plateau for the higher sizes of stimuli. Consequently, our estimation of the CMF will be lower for Monkey L.

**Figure 4.**
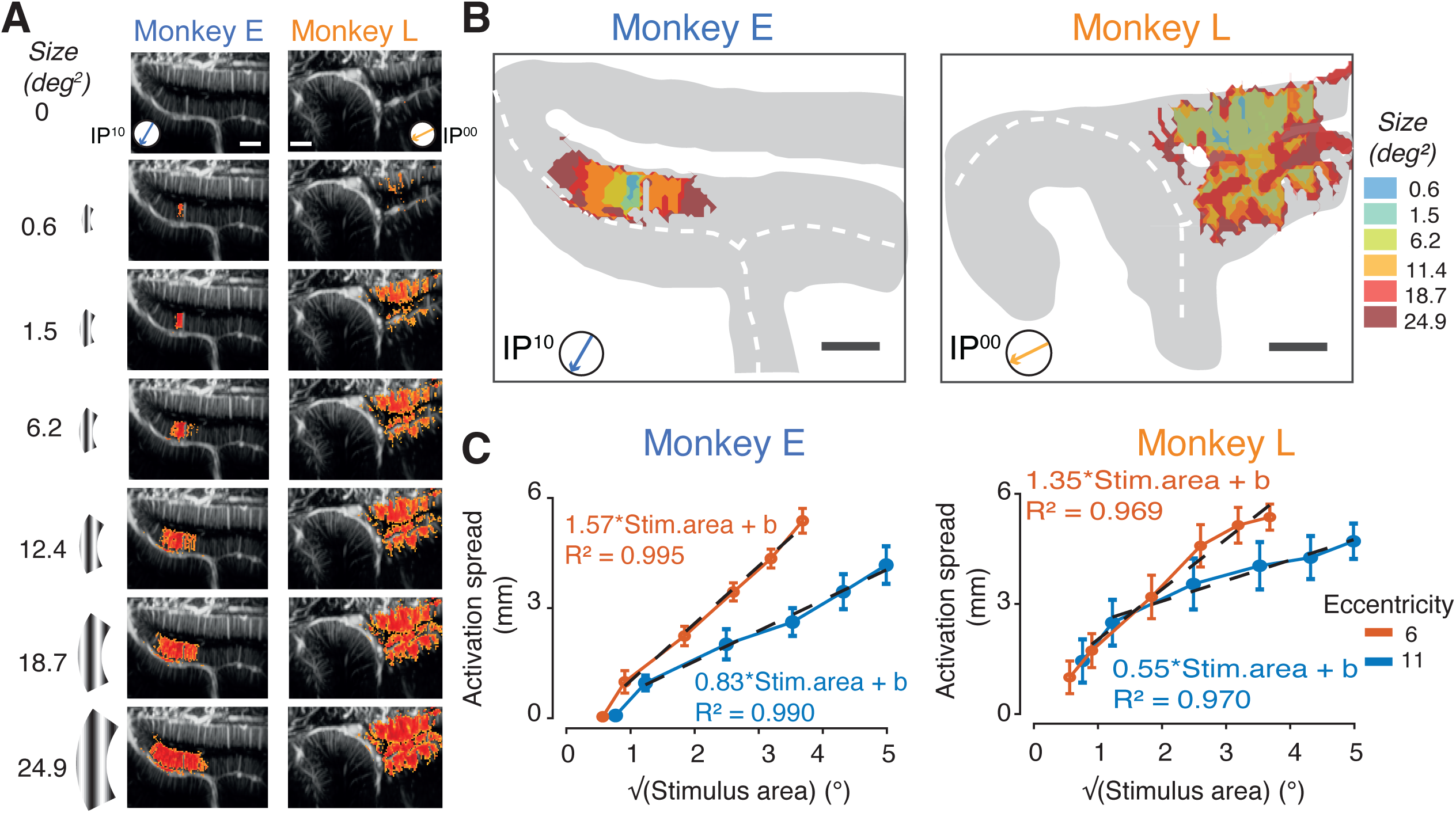
Effect of visual stimuli size on cortical activity. **(A)** Activation maps for different sizes of visual stimuli. Top representations: we designed 6 different stimuli to test the relation between the sizes of the visual stimulus and the superficies of the activation area; thickness is indicated in black (in DVA) and the value of opening angle in red (in deg). the first line of elements represents the different sizes tested for one location (polar position; *theta* = 213° and *rho* = 11 DVA). The size of the area (in deg²) varies according to the eccentricity (*rho*, with smaller values for an eccentricity if 6 DVA). For one condition, no stimulus was presented (No stim.). The 7 fUSi activation maps for a location (*theta* = 213 and *rho* = 11) of significant voxels (p = 0.001) per condition averaged on multiple sessions (n=5) for Monkey E (left column) and for Monkey L (right column). **(B)** Superposed activation maps. Color code indicate the size of the visual stimulus (deg^2^). **(C)** Evolution of the cortical spread activated as a function of the size of the visual stimulus (on the left Monkey E and the right panel for Monkey L). Eccentricities are indicated by the color code. Several locations were pulled together for the same eccentricity. The dashed lines represent the result of a linear regression computed to estimate CMF and compare these values to literature values. Error bars indicate ± SEM. Scale bars: 2 mm.

To benchmark our results, we compared them with established models of CMF. The original work introduced a linear magnification factor model (*31*). Later studies proposed a more accurate *LMF* including anisotropy of the factor in 2 directions: radial and angular. Both factors are described by the same power law with different coefficients:

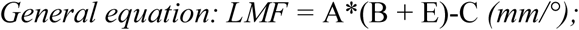

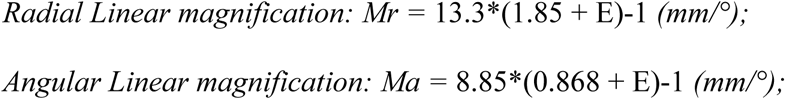

*with A* defined as the generalized scaling constant, *B* the generalized eccentricity at which the cortical representation is halved, *C* the precise exponent computed by regression and *E* the eccentricity. Using this theoretical model, we obtained values of *Mr* = 1.69 mm/deg and *Ma* = 1.28 mm/deg for the eccentricity of 6 DVA and *Mr* = 1.03 mm/deg and *Ma* = 0.74 mm/deg for the eccentricity of 11 DVA. We then compared these predictions to our experimental estimates by fitting cortical spread versus the square root of the stimulus area using linear regression. For both animals, we obtained significant fitting with R^2^ > 0.969 for Monkey L and R^2^ > 0.990 for Monkey E: *LMF* values were equal to 1.57 and 1.35 mm/deg at 6 DVA for Monkey E and Monkey L, respectively, and 0.83 and 0.55 mm/deg at 11 DVA. We assumed our measured LMF should be in between *Mr* and *Ma* for a given eccentricity (see discussion). Accordingly, all but one value respected this hypothesis, and we obtained values close to the theoretical values matching CMF studies done with vasculature and MRI data.

We then estimated the experimental value of the cortical point image. We first assumed that the point image remains constant across eccentricities, as proposed by the original study in fMRI on humans (*17*). We then defined the point image as the intersection of our two linear regressions fit (one for each eccentricity). Although this intersection is not directly visible in Figure 4C—due to the sensitivity limits of fUSi, which lead to reduced activity in the smallest condition— we extrapolated the regressions toward low stimuli sizes. This yielded an estimated point image value of approximately 0.4 mm for Monkey E and 2.82 mm for Monkey L due to a higher nonselective activity.

### Estimation of point image vasculature

In addition, we recorded Ultrasound Localization Microscopy (ULM) images on the same imaging planes used for fUSi to achieve a tenfold increase in resolution (from 100×100×400µm voxels for fUSi to 10×10×400µm voxels for ULM) for the anatomical images. We then superposed these high spatial resolution anatomical images (ULM) to the functional maps previously obtained by fUSi. This co-registering multimodal approach allowed us to test how stimulus size influences vascular activation within V1 and estimate the equivalent vasculature wise of the point image.

We acquired ultrasound localization microscopy (ULM) images in 12 planes from the same two anesthetized non-human primates (Monkey L and Monkey E see Figure S6). We selected the plane in which we recorded our fUSi data for the varying-stimulus-size task and co-registered the resulting vascular maps with the activation maps (Fig. 5A-D; Supplementary Fig. S5). These high spatial resolution images were obtained after microbubble intravenous injection (IV) under anesthesia; we were able to measure the flow velocity by tracking individual bubbles and reveal precise vascular mapping. Such an approach offered a detailed view of vascular organization within cortical folds, with higher resolution in 2D (10×10×400 µm voxels) than previous fUSi images (Fig. 1). In addition, by comparing the microbubble flow directions, we were able to differentiate arterioles from venules.

**Figure 5.**
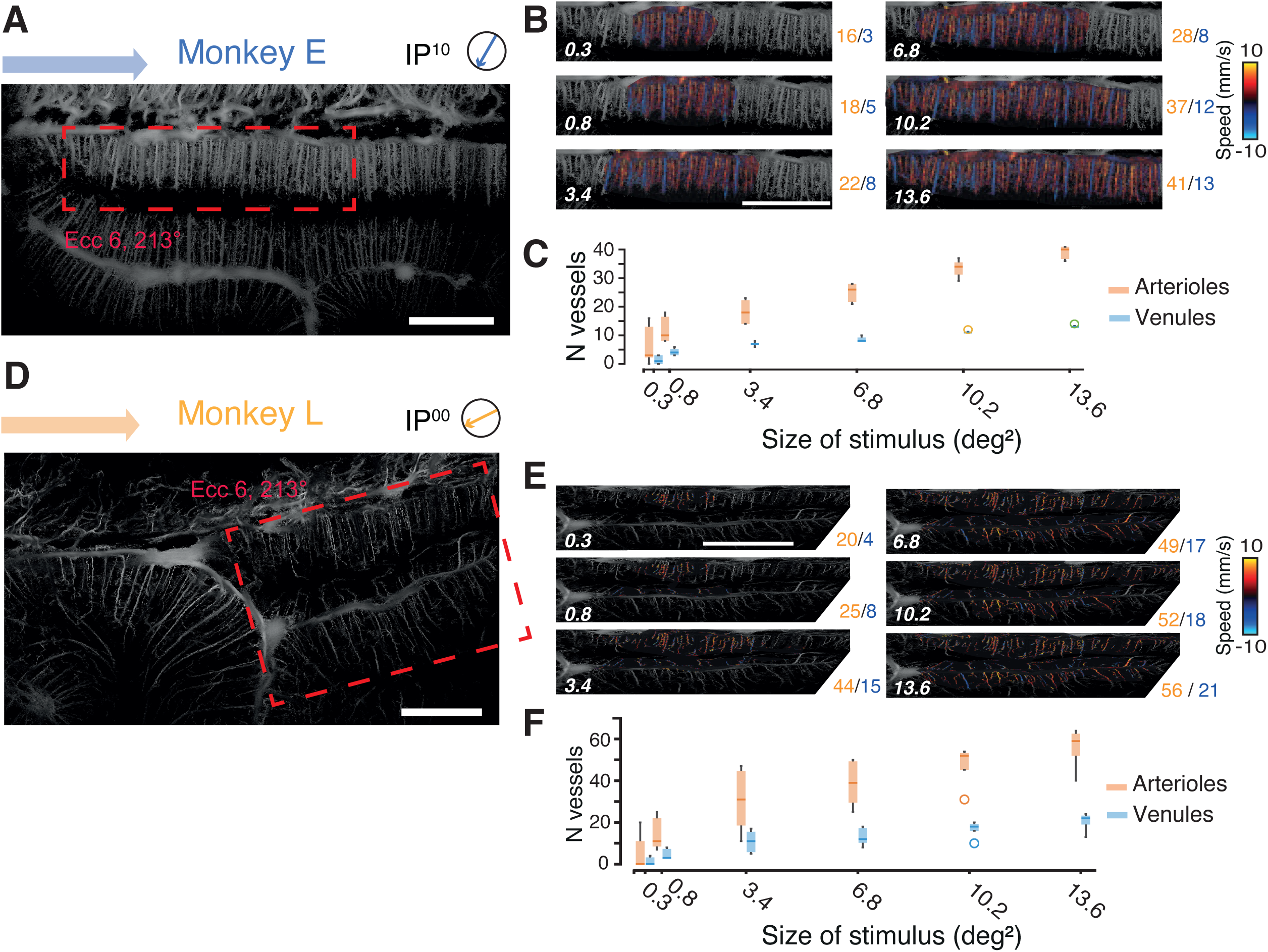
Vascular analysis combining functional ultrasound and ultrasound localization microscopy in an activated region. **(A)** ULM map for imaging plane IP^10^ for Monkey E generated from the counting of injected microbubbles, acquired at a resolution of 10 × 10 × 400 µm. The region of interest (ROI) for the one-stimulus position eccentricity 6 DVA, theta 213° is indicated in red. **(B)** Vessel counting on one example session: Number of arterioles (orange) and venules (blues). The white numbers indicate the size of the stimulus. **(C)** Number of arterioles and venules averaged on all sessions within the activity area for different stimulus sizes. **(D)** ULM map for imaging plane IP^00^ for Monkey L generated from the counting of injected microbubbles, acquired at a resolution of 10 × 10 × 400 µm. The region of interest (ROI) for the one-stimulus position eccentricity 6 DVA, theta 213° is indicated in red. **(E)** Vessel counting on one example session: Number of arterioles (orange) and venules (blues). **(F)** Number of arterioles and venules averaged on all sessions within the activity area for different stimulus sizes. Scale bars: 2 mm.

The vasculature density for Monkey E was consistent with anatomical reports in the literature (*3*) in the first fold (see supplementary table 3). We then overlapped fUSi activation maps for the visual stimuli presented at eccentricity 6 DVA with significant pixels located in this first fold. We thus counted the vascular structures (arterioles and venules) for each fUSi activation map as shown in Figure 5. For example, for Monkey E (Figure 5A), a stimulus size of 6.79 deg² at eccentricity 6 DVA, theta 191°, we had an activated area of 4.04 mm² (yellow area), that corresponds vasculature-wise to 23 arterioles and 9 venules recruited. Finally, the number of arterioles and venules recruited for all sizes of stimuli at eccentricity 6 DVA and for two different angles (theta = 191° and 213°) were computed and averaged on both animals (Fig. 5C-F). The number of vessels ranged from 0 to 19 for the lowest size of stimuli (0.3 deg², n=10) (Fig. 5A, right panel) indicating a high variability of inter-sessions with an average of 6.8 ± 7.6 vessels. From the other stimuli sizes, the standard deviation remained between ± 3.2 and ± 4.4 vessels (supp. Table 4), that were on the order of magnitude of arterioles/venules ratio for superficial layers for both animals (supp Table 3). Except for very small stimulation sizes, close to the fUSi sensitivity limits, the small standard deviation demonstrated the reproducibility of the approach. Similarly, to Monkey E, we analyzed responses to stimuli presented at eccentricity 6 DVA (Figure 5B) for Monkey L. For such stimuli, the mean number of vessels (see supp. table 4) identified was higher due to a strong activity in the first fold present for each size. This correlates once more with the non-selectivity of this fold brought to light with the retinotopy standard deviations.

Using the previously established vessel density (*3*) and point image values, we can estimate the number of vessels contained within the functional point image structure across our 400µm thick imaging plane. In Monkey E, the point image surface of 0.16 mm² corresponds to an average of 1.9 arterioles and 0.8 venules, while in Monkey L, the larger point image surface of 1.12 mm² corresponds to an average of 13.5 arterioles and 5.6 venules.

## Discussion

The present study demonstrates that using functional ultrasound imaging (fUSi) enables high-resolution, large-volume mapping of macaque primary visual cortex across cortical folds and layers. Moreover, by co-registering fUSi with ultrasound localization microscopy (ULM) we were able to link point image volume and cortical vasculature. A limited number of sessions were sufficient to obtain stable, reproducible retinotopic maps with enough resolution to study its dependencies from infragranular to supragranular cortical layers and to map the variations of cortical activation spread in response to different sizes of stimuli. We were also able to derive from those maps a cortical magnification factor (CMF) consistent with prior literature and challenge the latest estimations of the point image as a functional unit. Finally, ULM provided complementary vascular detail, allowing us to relate functional patches to underlying microvasculature and to quantify arteriole/venule engagement as a function of stimulus size.

Using fUSi, we obtained clear and distinct retinotopic maps across different regions of primary visual cortex along anatomical sulci as lunate and calcarine. In Monkey E, the maps revealed a systematic progression of preferred eccentricity values that closely followed the cortical geometry of the calcarine sulcus, with eccentricity preferences organizing into contiguous bands that tracked each successive fold. A very similar organization has been reported with macaque fMRI retinotopic mapping using phase-encoded expanding rings (eccentricity) and rotating wedges (polar angle)(*28*, *29*, *33*). In contrast, Monkey L exhibited a less coherent organization in both eccentricity and polar-angle representations across imaging planes. We have noticed in our fUSi maps a variability across-sessions comparable with the ones reported with fMRI. For example, on human fMRI, polar-angle mean absolute errors typically lie in the low–mid tens of degrees (≈ 25 ± 11DVA) from a single short scan, while eccentricity errors are sub-degree (0.76° ± 0.34°) ∼0.41° median, (*34*, *35*), with reliability declining toward the stimulus edge (*36*). In our study, we reported a variance of the preferred angle of 21° and 42° for respectively Monkey E and Monkey L and a variance for eccentricity of 2.0 and 2.6 DVA for respectively Monkey E and Monkey L (see Figure S4).

The fine-grained laminar organization —of V1 observed in literature (*37–39*) echoes the laminar trends observed in our data. Our layer-resolved analyses showed that retinotopic preferences were largely conserved across laminae for supragranular, granular and infra granular layers, with modest depth-dependent shifts in preferred eccentricity and response amplitude. Supragranular ensembles tended to exhibit slightly broader activations than infragranular ensembles, with the granular band most confined—patterns consistent with thalamocortical input to L4 and known depth variations in microvascular density. However, we could also be missing the majority to blood influx variation between layers as we were not able to image capillaries with the actual fUSi technique. As blood is provided to arterioles that cross all layers, we have fewer chances of detecting the difference between layers. A hint to the variations visible through our imaging could be a small but detectable variation of vessel diameter before sending blood to the capillaries. fUSi offers a distinct compromise: ∼100-µm resolution allowing deep and continuous imaging of cortical folds through a chamber, with better coverage than optical methods and less invasiveness that electrode arrays (*40*). Notably, the spatial sensitivity of fUSi significantly exceeds that of ultra-high field 7T fMRI laminar protocols, which are generally restricted to 500– 800 µm voxels (*15*). In human fMRI, related projections are typically derived from pRF models and summarized on cortical surfaces to delineate map borders and field coverage (*34*, *41*). By contrast, optical approaches (IOS/VSDI) achieve high spatial resolution for projecting fine retinotopic structure, yet are restricted to gyral crowns and superficial sulcal lips, leaving the calcarine and lunate fundi under-sampled in vivo (*12*, *42*).

Classically, the CMF values are decreasing with eccentricities, reflecting the stronger representation of central vision in V1(*31*, *32*). We similarly observed that activated area was larger at lower eccentricities (approximately 1.5 times more activity at eccentricity 6 compared to eccentricity 11 for a 14 deg² visual stimulus). Our experimental CMF values were aligned with the theoretical benchmarks of Horton & Hoyt (1991) and Adams & Horton (2003). These models used vasculature to compute Linear magnification factor (LMF) and account for anisotropy by having two LMF (*32*), one for the radial dimension of the visual field (Mr) and one for the angular dimension of the visual field (Ma). Because fUSi was performed in 2D, we captured activity spread along a single dimension, assuming to lie between the radial and angular components. Indeed, theoretical values at 6 DVA were *Mr* = 1.69 mm/deg and *Ma* = 1.28 mm/deg and our estimates are 1.57 and 1.35 mm/deg for Monkey E and Monkey L respectively and at 11 DVA *Mr* = 1.03 mm/deg and *Ma* = 0.74 mm/deg while our estimates are 0.83 and 0.55 mm/deg for Monkey E and Monkey L respectively. The hypothesis was not valid for one condition, for Monkey L at 11 DVA, because of the presence of non-selective activity in the superficial fold, especially for the small sizes of stimuli. When we compared our results to previous approaches, our results extend the scope of electrophysiological, optical imaging (*43–46*) and fMRI work by showing that mesoscale hemodynamic imaging can provide subject-level CMF estimates in non-human primates. Then, the point image values obtained in the present study differed substantially between subjects, measuring approximately 0.4 mm for Monkey E and 2.82 mm for Monkey L. Notably, for Monkey L, correcting for non-selective neural activity would be expected to yield a point image estimate more closely aligned with that of Monkey E. Taken together, these findings converge toward values lower than those previously reporting (*17*) suggesting that earlier fMRI-based estimates may have been overestimated by the relatively coarse fMRI spatial resolution available at the time. Altogether, these results indicate that fUSi can resolve functional activation at a scale finer than previously accessible with non-invasive imaging, while remaining broadly consistent with the mesoscale hemodynamic framework established by prior CMF analyses.

Histological studies have defined vascular densities in primate V1 (*3*). We here report the vascular densities in living primates throughout the whole cortical layers. This investigation successfully demonstrated the utility of ULM to describe the link between mesoscopic cortical activity observed with fUSi and the underlying microvasculature network (*24*) in non-human primates. The super-resolution functional imaging modality has primarily been validated in rodent models (*22*, *23*, *47*) and more recently in humans through the temporal bone, enabling resolutions up to 25µm (*48*). Our implementation builds upon these studies by co-registering anatomical ULM with fUSi. We were able to quantify the recruitment of arterioles and venules in a manner consistent with the histological densities reported in macaque V1 (*49*) while the high variability in vessel counts for the smallest stimuli (0.3 deg²) underscores a clear sensitivity floor.

In summary, the integration of ULM provides a microscopic anatomical framework that clarifies the structural underpinnings of the fUSi signal, even as it highlights the current technical boundaries of functional ultrasound. Although fUSi can detect activity-driven hemodynamic changes at a mesoscopic scale, defining a singular discrete “fUS vascular unit” remains challenging due to the stochastic nature of microvascular recruitment near the limits of its sensitivity. Despite these limitations, this multi-scale approach effectively addresses a critical bottleneck in high-resolution neuroscience: the difficulty in distinguishing between specific microvascular perfusion and non-specific macrovascular drainage. While even ultra-high-field 7T fMRI remains largely biased toward the BOLD signal of larger pial veins, the ability of ULM to isolate the flow dynamics of individual microvessels provides a structural ground truth. This establishes fUSi co-registered with ULM anatomical maps as a first step for investigating the 3D functional architecture of the primate brain, offering a level of microvascular specificity that remains difficult to achieve with non-invasive MRI protocols

To conclude, our results on CMF and ULM also raise several perspectives. First, beyond 2D mapping, 3D fUSi could enhance precision and fast determination of each subject CMFs and the extension to 3D volumetric functional ULM could achieve near-isotropic resolution (e.g., 40×40×40 µm), potentially resolving single arteriole-venule pairs and providing a comprehensive 3D atlas of neurovascular coupling. Second, from a translational perspective, estimating subject-specific CMF with fUSi could help design and calibrate cortical stimulation protocols to produce visual percepts and the ability of functional ULM to non-invasively map the mesoscopic human brain through acoustic windows could provide a new window into neurovascular disorders, where early microvascular changes often serve as the first biomarkers of disease. Finally, combining fUSi with higher-resolution ULM opens the possibility of linking vascular units to the minimal resolvable stimulus sizes, thereby bridging scales from population-level point images to single-vessel dynamics.

## Materials and Methods

### Behavioral training

We collected data from two non-human primates: a female rhesus macaque (“Monkey L”; 14 years, 10 kg) and a male cynomolgus macaque (“Monkey E”; 14 years, 7 kg). Animals were individually housed and cared for in accordance with institutional Welfare Committee guidelines on good animal practice; all procedures were authorized by the French Ministère de l’Éducation nationale, de l’Enseignement supérieur et de la Recherche (APAFIS #9013-2017021515254591) and approved by the institutional and regional animal care committees (Comité C. Darwin). During experiments, monkeys were seated in a primate chair (Crist Instruments), head-fixed, and positioned 58 cm from a cathode-ray (CRT) computer screen in a darkened booth; mean screen luminance was maintained at 1.15 mW·cm⁻². Eye position was recorded at 1 kHz with an infrared video eye tracker (EyeLink 1000, SR Research), enabling online control of the behavioral paradigm and delivery of a liquid reward (sucrose solution) contingent on task performance, with experimental control implemented in EventIDE (Okazolab, Netherlands). Primates were maintained under mild fluid restriction (≈30 mL·kg⁻¹·day⁻¹) and were allowed to drink ad libitum while working.

### Visual Stimuli paradigm

Monkeys were trained on a passive fixation task. Each trial began when the animal fixated a central green square (0.2° visual angle) for 1 s within a 2.5° circular window; a peripheral stimulus was then presented in the left visual field for 1 s. For eccentricity mapping, we used 2 DVA-wide hemiconcentric bands centered on fixation, presented at eccentricities from 0 DVA (catch) to 14 DVA in 2 DVA increments. For polar-angle mapping, we used 15°-wide wedges spanning 1.5–15 DVA eccentricity, centered at polar-angle positions randomly chosen between 7.5° and 182.5°. To test different sizes of stimuli we used a portion of an annulus and increased the angular opening and thickness of the annulus simultaneously. All stimuli consisted of sinusoidal gratings (spatial frequency: 1 cycle/deg) oriented horizontally presented with a fixed temporal frequency (2.78Hz). Correct trials were rewarded with a small drop of sucrose solution, and stimulus onsets were separated by 10 s to allow cerebral blood volume to return to baseline.

### Functional ultrasound imaging (fUSi) recordings

The head was stabilized with a surgically implanted titanium head post (Crist Instruments, MD, USA). After behavioral training, we implanted a CILUX recording chamber (Crist Instruments, MD, USA) and performed a 19-mm craniotomy centered for Monkey E at the coordinates mediolateral +7 mm, anteroposterior −22 mm and for Monkey L: mediolateral +7 mm, anteroposterior −10 mm). A custom 15-MHz linear ultrasonic probe (128 elements; in-plane spatial resolution 100 × 100 μm; slice thickness 400 μm) was used in the chamber with sterile ultrasound gel. This configuration provided a field of view spanning ∼12 mm along the cortical surface and up to 15 mm in depth. Before each session, the chamber was filled with sterile gel (Aquasonic 100, Parker Laboratories), and the probe was mounted on a custom adapter and positioned perpendicular to the cortical surface, making gentle contact through the gel.

Changes in cerebral blood volume were recorded with a real-time functional ultrasound scanner prototype (Iconeus and Inserm U1273, Paris, France) using the custom 15-MHz probe. Data was acquired in continuous bursts of 11 plane-wave transmissions spanning angles from −10° to +10° (pulse-repetition frequency, PRF: 5.5 kHz). Echoes from each burst were coherently summed to form a compound image every 2 ms (effective frame rate: 500 Hz). Power Doppler images were then obtained at 2.5 Hz by averaging 200 consecutive compound frames following spatiotemporal clutter suppression based on singular value decomposition (SVD). We only selected correct trials for fUSi analysis.

### Ultrasound Localized Microscopy recordings

ULM acquisitions were performed across twelve distinct imaging planes per primate. For each plane, we acquired a 3-min contrast-enhanced ultrasound recording optimized for microbubble tracking under isoflurane anesthesia. After a 30-s baseline, 1 mL of SonoVue was injected intravenously, followed 1 min later by a second 1 mL bolus; acquisition continued throughout to complete the 3-min recording. After each plane, the probe was advanced to the adjacent plane, and all twelve planes were recorded sequentially within a single session for each animal.

## Data Processing

### · Calculation of mean retinotopic map and variability across sessions

For both eccentricity and polar-angle mapping, the final retinotopic maps were obtained by first computing a session-specific winner-takes-all map and then combining these maps across repeated sessions. Specifically, for each session and each voxel, we estimated the response to every stimulus condition (12 polar-angle wedges or 6 eccentricity arc-bands) and assigned to the voxel the stimulus label that elicited the strongest response (maximum Z-score). This yielded, for each session, a discrete retinotopic map in which every voxel was associated with a single preferred polar angle (wedge center angle) or preferred eccentricity (band center eccentricity).

To obtain the final maps shown in the main figures, we spatially aligned all session maps to a common anatomical reference within each imaging plane and then computed, voxel-by-voxel, the central tendency of the preferred values across sessions (i.e., the mean/consensus preferred polar angle or eccentricity after alignment).

We quantified the across-session variability of retinotopic preference at the voxel level as the dispersion of the session-specific preferred values. For each voxel, we collected the set of preferred polar angles (or eccentricities) assigned across all sessions and computed the standard deviation of these values. This resulted in a variability map in which higher values indicate voxels whose preferred stimulus label fluctuated strongly across sessions, whereas lower values indicate stable retinotopic assignment.

### · GLM analysis

Doppler data were analyzed with a parallelized generalized linear model implemented in MATLAB (MathWorks). For each condition, the inputs were the Doppler time series and the corresponding stimulus timestamps. A delta-function train (Dirac comb) marking stimulus onsets was convolved with the hemodynamic response function to generate a condition-specific CBV predictor. For every voxel, the GLM fit yielded a z-score and p-value for the condition regressor; activation maps display z-scores for voxels surviving p < 0.001 after Benjamini–Hochberg false-discovery-rate correction. Retinotopic maps for a single acquisition were obtained by assigning to each voxel the condition producing the maximum z-score across the eccentricity or polar-angle set, and maps from repeated runs within the same imaging plane were averaged to yield mean preferred-eccentricity and preferred-polar-angle maps.

### · ROI analysis

After GLM estimation, we restricted multiple-comparison correction to a manually defined region of interest (ROI) to limit the number of tested voxels. Specifically, the Benjamini–Hochberg false-discovery-rate procedure was applied within the ROI, reducing the correction burden and increasing sensitivity (i.e., retaining more truly active voxels) while maintaining FDR control. ROIs were drawn around the calcarine sulcus for Monkey E and around the lunate sulcus for Monkey L.

### · Image registration between sessions

At the start of data collection, we acquired a reference anatomical image for each imaging plane in both Monkey L and Monkey E. All subsequent sessions from the same plane were aligned to this reference to correct occasional global frame shifts observed across acquisitions—shifts attributable to transient changes at the probe–tissue interface caused by rare, forceful movements. Corrections were performed with 2-D rigid-body image registration (translation + rotation) using MATLAB’s imregcorr.

### · Cortical layer specification

To parcellate the cortical fold nearest the surface into laminar ensembles, we first manually delineated the cortical surface (border between cortex and adjacent non-neural tissues) on the mean eccentricity and polar-angle maps. Voxels located 0–500 µm below this surface (i.e., 0–5 voxels at ∼100 µm/voxel) were assigned to layers I–III; voxels 500–1100 µm deep were assigned to layer IV; and voxels 1100–1500 µm deep were assigned to layers V–VI. These depth bins follow published thickness estimates for macaque V1. We applied an analogous procedure to the deeper cortical fold, noting that the laminar order is reversed along the dorsoventral axis: we first delineated the ventral cortical border and then extracted the same depth bands dorsally to define layers I–III, IV, and V–VI.

### · Stimuli Size analysis

We used the GLM analysis described earlier with a modified Dirac comb for the Stim size test. Since all visual stimuli are centered on the same position, many voxels are activated by multiple conditions. Therefore, we adapted the Dirac combs for each condition. For example, for the “N”th one size of the stimulus, we created N combs. The first comb includes the timings of all the conditions visually smaller than N, the second comb all the conditions smaller than N without the smallest, and so on. The last comb would be the timings of the condition N corresponding to the voxel specific to the condition displaying the highest Z-score from those previous GLM analyses for each voxel of this condition. Without this processing, we observed, for more prominent visual stimuli, a decorrelation between the power Doppler signal and activity of the voxels at the center of the activated cortical area due to constant activation.

For the linear regression we included data points starting from the second size of stimuli to avoid the non-linearities present at small sizes of stimuli (*32*). As we can see in both primates the first data point drops significantly in a nonlinear manner compared to the rest of the curve.

### · Ultrasound localization microscopy image reconstruction

Raw functional ultrasound data blocks consisting of 200 frames were initially processed by separating tissue signals from those of microbubbles, achieved by removing the first twenty components of the Singular Value Decomposition. Each frame was subsequently sampled sixfold using Lanczos interpolation. Microbubbles were detected as the brightest local maxima in the correlation with a typical point spread function (Gaussian spot). Only those microbubbles with a correlation value exceeding 0.7 were retained. A second-order spatial polynomial fit was applied to determine the sub-voxel peak location corresponding to the center of each individual microbubble which was then projected onto a [6.5 µm × 6.5 µm] grid. A particle tracking algorithm (https://github.com/tivenez/simpletracker) was employed to track the maxima positions at each time step and successive positions of individual microbubbles were compared to the previous ones to derive velocity vector fields. Maps of microbubble count, velocity and back-scattering amplitude were generated by accumulating all detected microbubbles and their amplitude within each voxel.

### · Quantification of vessel population in the visual area stimulated

ULM images were reconstructed in the plane of the 3D rotational scan that matched the one used during visual stimulation experiments. The two corresponding power Doppler images were aligned using a rigid geometric transformation, implemented with the ’imregtform’ function in MATLAB. For each visual stimulus, a polygonal region of interest was manually drawn around the activated area on the activation map. Vessels were identified by applying a vessel-enhancing filter (Vesselness 2D) to the backscattering images, retaining only structures with an ovoid shape and a size greater than 200 voxels. Finally, the spatially averaged microbubble velocity along the depth axis was calculated for each vessel. Vessels were classified based on flow direction: if the velocity was directed toward a large horizontal vessel, it was identified as a vein; if the flow was directed away, it was classified as an artery.

### · Visual field and anatomical visual projection

To realize our projections, we used the mean averaged retinotopic map previously produced. For the voxel of our anatomical doppler image that had a non-null value in BOTH mean eccentricity map and mean phase map we associated their preferred eccentricity and phase to them which gave us their position in the visual field. The color code indicated in which cortical layer the position within the visual field was encoded.

## Acknowledgments

Part of this work was carried out at the [Paris Brain Institute PHENOPRIMR Core Facility](https://rrid.site/data/record/nlx_144509-1/RRID:SCR_028519/resolver?q=phenoprimr&i=rrid:scr_028519) (RRID:SCR_028519). We gratefully acknowledge Morgane Weissenburger, Estelle Chavret-Reculon for day-to-day management of the animal facility and Bénédicte Daboval and Lucile Aubrée for veterinary care. We warmly thank Chris Klink for his careful reading of the manuscript and for his insightful comments and correction.

## Author contributions

Conceptualization: FA, ED, IA, MP

Methodology: ED, IA, MP, RA

Investigation: ED, IA, MP, RA

Visualization: ED, IA, MP, NZ, FA

Supervision: FA, PP, SP, MT

Writing—original draft: ED, IA, MP, FA

Writing—review & editing: FA, FF, PP

## Data and materials availability

All data. Code and materials reported in this paper are available in the Open Sciences Framework

## Supplementary Materials

**Figure S1 All fUSi anatomical planes for both animals Monkey E and Monkey L. (A)** 12 fUSi imaging planes (IP00 to IP55). The white circle represents within each panel the recording chamber and the colored arrow (orange for Monkey L and bleu for Monkey E) the position of the fUSi probe. The head of the colored arrow on the top of the image indicates the orientation of the fUSi images. Standard orientations of the image are indicated (A: anterior, P: posterior, M: Medial and L: Lateral). (**B**) Same representations as in A but for Monkey L. Scale bars : 2 mm.

**Figure S2 Retinotopic maps based on different eccentricities. (A)** Activation maps according to the 6 different eccentricity positions of the visual stimulus (here a semi-circular portion of a circular vertical grating). Same representation as in Figure 2. Each image represents the activation map for a particular eccentricity angle (2 to 14, each 2 DVA). The color code indicates the value of the z-score. **(B)** A single session eccentricity map. Eccentricity map reconstructed from activations in A, obtained in a single session for 6 different eccentricity stimulations (∼20 trials per eccentricity angle). **(C)** Mean eccentricity map for one imaging plane (IP10). We averaged the total number of sessions (n=8) to obtain eccentricity maps for one specific imaging plan. **(D and E)** Mean eccentricity maps for other imaging planes for Monkey E (D) and for Monkey L **(E)** respectively. Same representation as in Figure 2. Scale bars: 2 mm.

**Figure S3 Averaged polar angle and eccentricity retinotopic maps for all fUSi planes (IP00 to IP55). (A)** Averaged functional maps for polar angles. The 12 top images are for Monkey E: (**B)** The 12 bottom ones for Monkey L. The color code indicates the preference for polar angle. Only all significant z-score voxels are represented. (**C)** Averaged functional maps for eccentricity. The 12 top images are for Monkey E. (**D)** The 12 bottom ones for Monkey L. The color code indicates the preference for eccentricity. Only all significant z-score voxels are represented. Scale bars: 2 mm.

**Figure S4: Standard error variation across all fUSi planes for polar angle and eccentricities. (A)** Each image represents for one fUSi (IP00 to IP55) the repartition of the standard deviation across the multiple sessions indicated by n. The 12 top images represent SD maps for polar angle for Monkey E, the 12 bottom ones for Monkey L. Color code indicates the value of the SD. Symbols are similar to Figure S1. **Right panel.** Representation of the averaged standard deviation. Each bar represents the average SD for one fUSi plane across the total number of sessions. The black bar represents the average SD across the total number of sessions and fUSi planes (n=12). **(C)** Same representation but for Monkey L**. (C)** Same representation as in panel A but for eccentricity for Monkey E. **(D)** Same representations as in panel B but for eccentricity for Monkey L. Scale bars : 2 mm.

**Figure S5 Single session data for CMF factor. (A)** Single session data representing the spread of cortical activity in function of the square root of the area of the visual stimulus for Monkey E (top panel, n=4) and 2 for Monkey L (bottom panel, n=5). The stimulus is centered at eccentricity 6 and at 3 different angles (solid, dotted and dashed lines). **(B)** Single session data representing the spread of cortical activity in function of the square root of the area of the visual stimulus for Monkey E (top panel, n=4) and 2 for Monkey L (bottom panel, n=5). The stimulus is centered at eccentricity 11 and at 2 different angles (solid and dashed lines). Error bars represent ± SEM.

**Figure S6 ULM anatomical images for all imaging planes. (A)** ULM images for the 12 used fUSi planes (IP00 to IP55) for Monkey E. **(B)** Same representations as in panel A but for Monkey L. Symbols are similar than in Figure S3. Scale bars : 2 mm.

**Table S1 Variations of Eccentricity between cortical layers.** Represents for both animals, for all significant pixels the mean retinotopic eccentricity value with standard deviation per cortical fold per layer. Complementarily, the p value of one-side paired T-test with Benjamini corrections by pairs of layers inside each fold.

**Table S2 Variations of Angle between cortical layers.** Represents for both animals, for all significant pixels the mean retinotopic angle value with standard deviation per cortical fold per layer. Complementarily, the p value of one-side paired T-test with Benjamini corrections by pairs of layers inside each fold.

**Table S3 Vessel density per cortical fold.** Represents for both animals, the mean (across all planes) arterioles and venules density (vessels/mm²) with standard deviation per cortical fold and their subsequent ratio.

**Table S4 Vessel counting per cortical fold.** Represents for both animals, the mean (across all planes) number arterioles and venules with standard deviation per cortical fold.

