## Supplementary figures and images for "From Functional Organization to Vascular Structure in the Primate Visual Cortex Using Ultrasound Imaging"

### Supplemental Figure 1

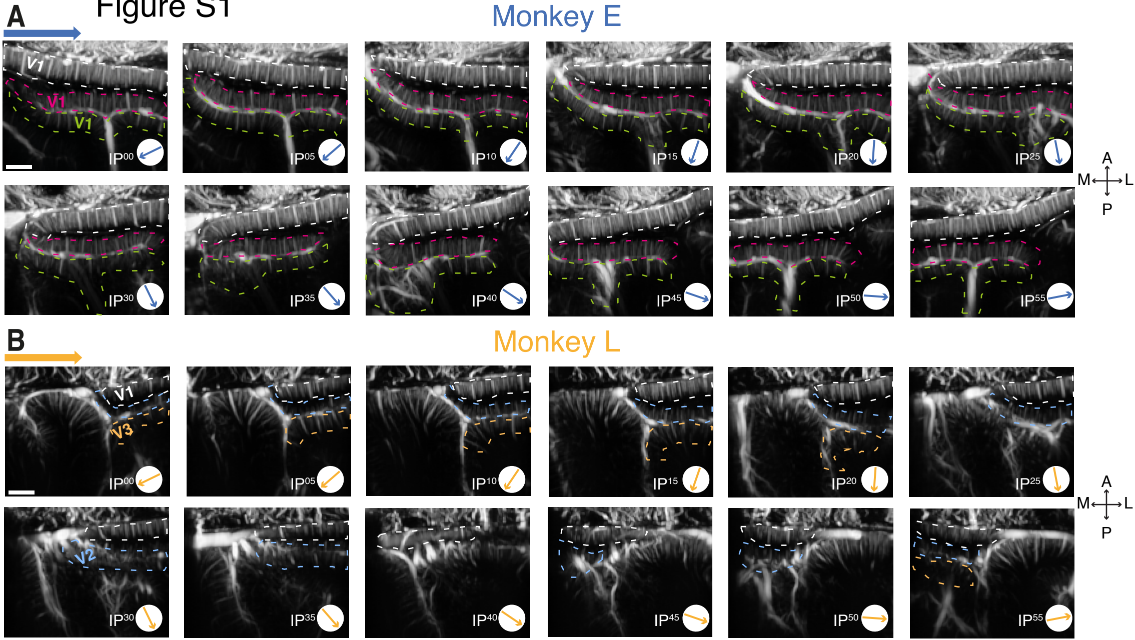

### Supplemental Figure 2

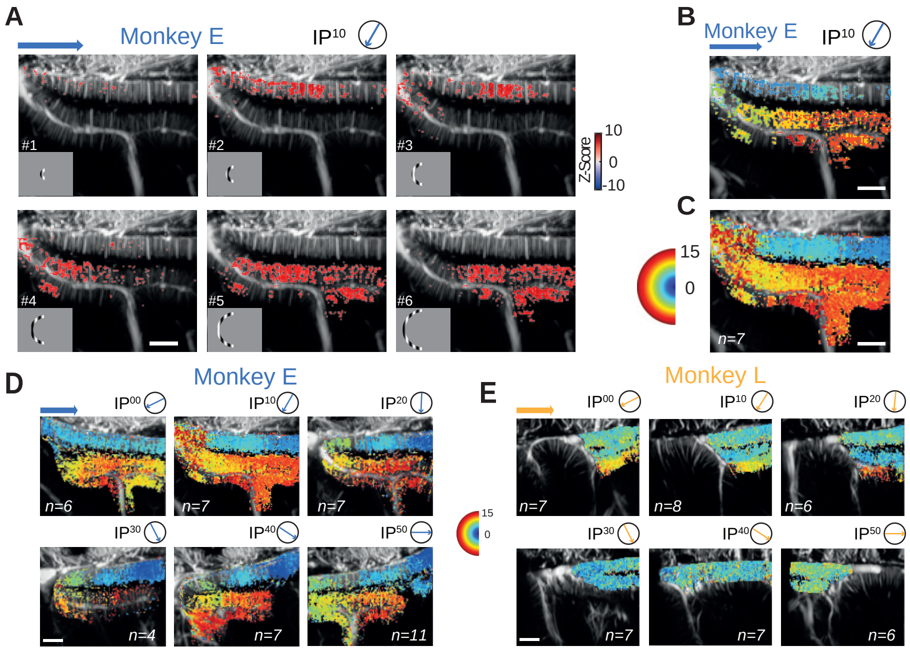

### Supplemental Figure 3

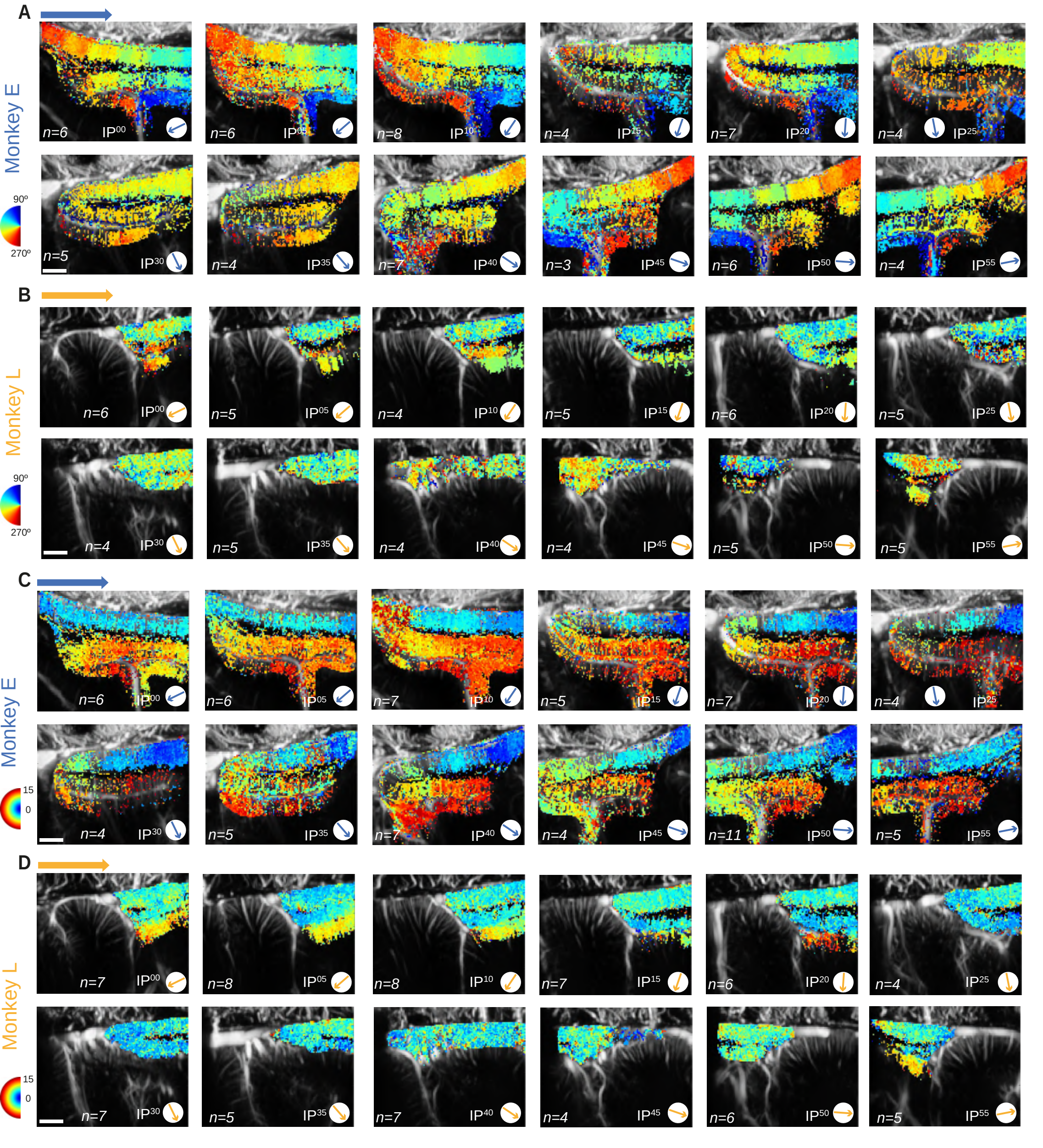

### Supplemental Figure 4

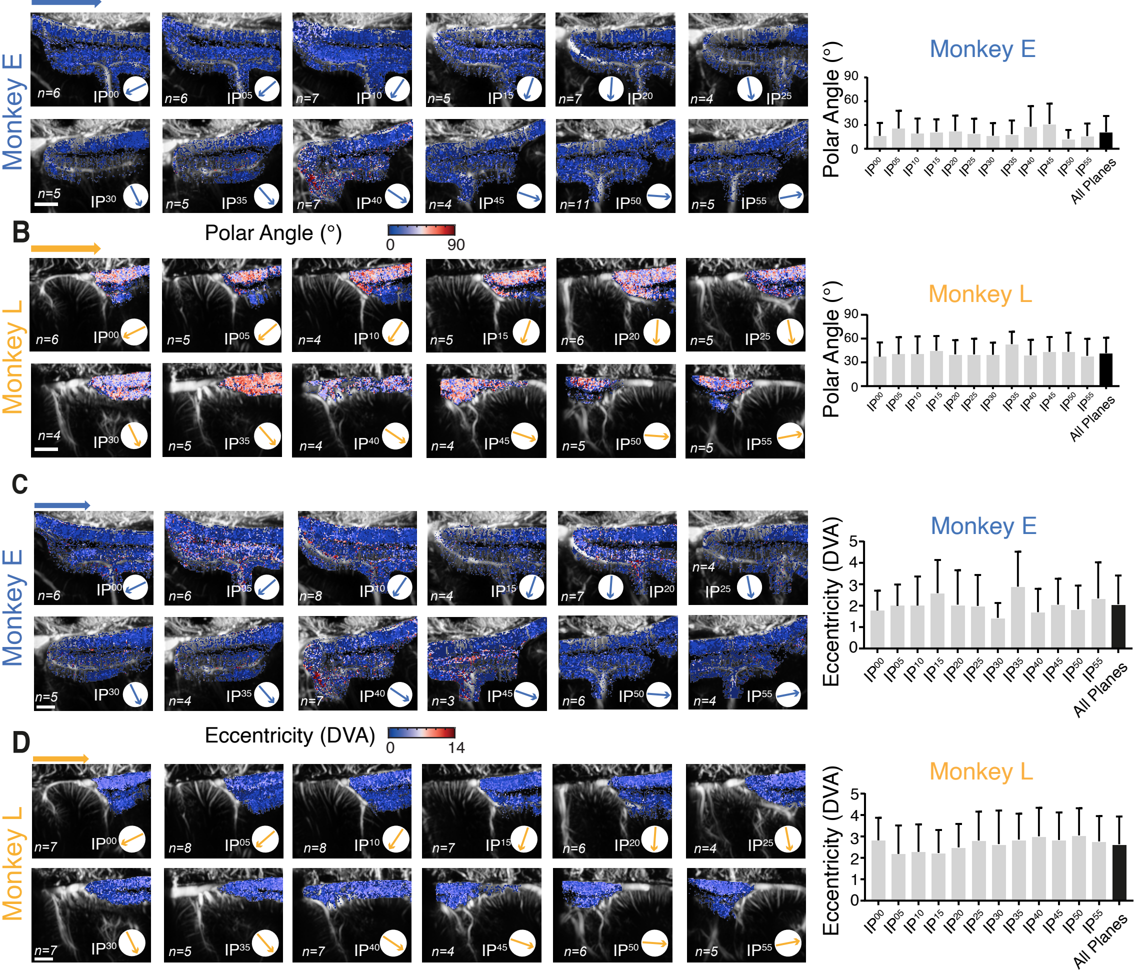

### Supplemental Figure 5

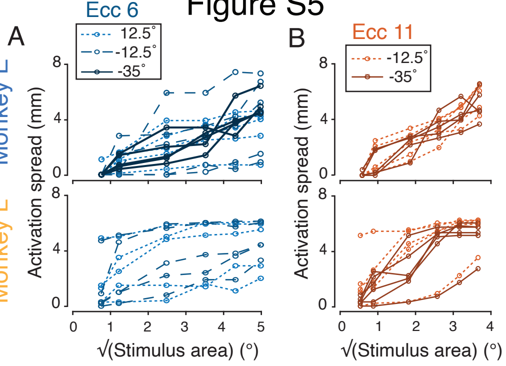

### Supplemental Figure 6

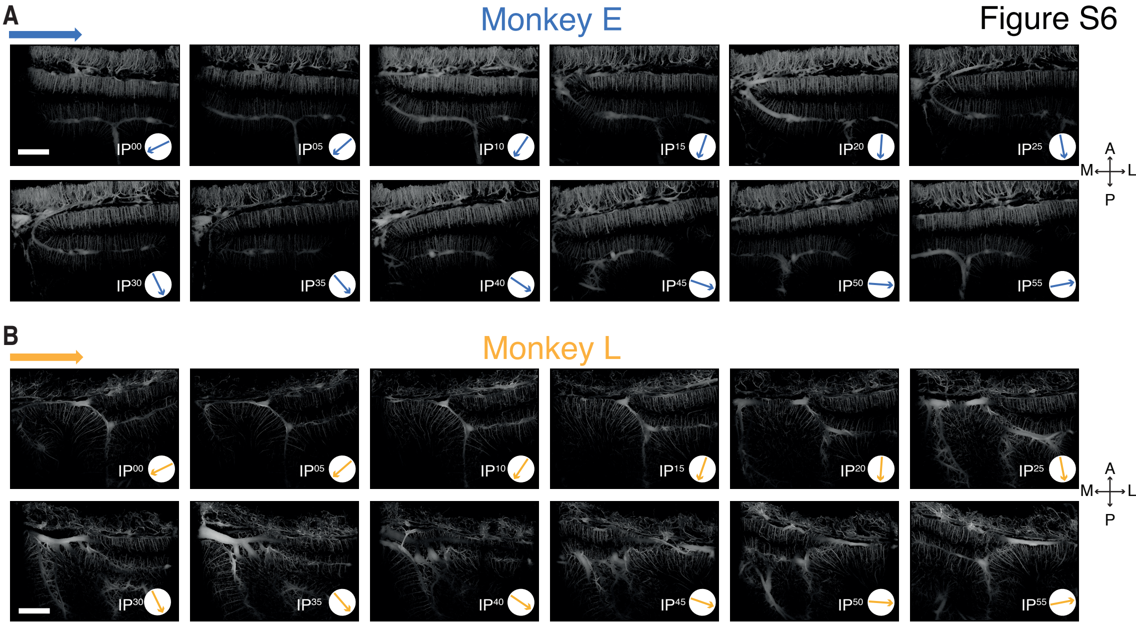
