## Supplemental Tables 1 and 2 for "From Functional Organization to Vascular Structure in the Primate Visual Cortex Using Ultrasound Imaging"

### 1. Descriptive values for Eccentricity (DVA)

| *Animal* | *Cortical fold* | *Layer* | *Mean preferred eccentricity* | *p value* | *N voxels* |
|---|---|---|---|---|---|
| Monkey E | V1 superior fold | L1 & L2/3 | 5.2 ± 1.9 | 3.07E-11 | 2137 |
| | | L4 | 5.3 ± 2.1 | | |
| | | L5 & L6 | 5.5 ± 2.2 | 9.03E-10 | |
| | V1 inferior fold | L1 & L2/3 | 9.5 ± 1.8 | 2.13E-11 | 1632 |
| | | L4 | 10 ± 1.8 | | |
| | | L5 & L6 | 10.2 ± 2.0 | 1.27E-20 | |
| Monkey L | V1 | L1 & L2/3 | 6.1 ± 0.9 | 3.35E-02 | 1396 |
| | | L4 | 6.1 ± 0.8 | | |
| | | L5 & L6 | 6.6 ± 1.2 | 2.69E-42 | |
| | V2 | L1 & L2/3 | 6.4 ± 1.3 | 1.20E-11 | 1217 |
| | | L4 | 5.7 ± 1.1 | | |
| | | L5 & L6 | 6.0 ± 1.5 | 1.55E-42 | |

*p values for one-side paired T-test with Benjamini corrections*

### 2. Descriptive values for Polar angle (º)

| *Animal* | *Cortical fold* | *Layer* | *Mean preferred angle* | *p value* | *N voxels* |
|---|---|---|---|---|---|
| Monkey E | V1 superior fold | L1 & L2/3 | 194 ± 24 | 1.98E-4 | 2669 |
| | | L4 | 195 ± 24 | | |
| | | L5 & L6 | 195 ± 23 | 2.27E-2 | |
| | V1 inferior fold | L1 & L2/3 | 191 ± 24 | 1.1E-3 | 1316 |
| | | L4 | 191 ± 22 | | |
| | | L5 & L6 | 190 ± 23 | 4.95E-1 | |
| Monkey L | V1 | L1 & L2/3 | 173 ± 17 | 3.5E-14 | 1256 |
| | | L4 | 176 ± 15 | | |
| | | L5 & L6 | 172 ± 19 | 1.75E-14 | |
| | V2 | L1 & L2/3 | 183 ± 22 | 2.86E-22 | 1000 |
| | | L4 | 175 ± 18 | | |
| | | L5 & L6 | 176 ± 16 | 1.1E-1 | |
