## Supplemental Tables 3 and 4 for "From Functional Organization to Vascular Structure in the Primate Visual Cortex Using Ultrasound Imaging"

### 3. Vessel density

| | *Cortical fold* | *Vessel* | *Mean density + std (vessels/mm2)* | *Ratio* |
|---|---|---|---|---|
| ***Monkey E*** | Superficial | arterioles | 12.8 ± 1.0 | 2.4 |
| | | venules | 5.3 ± 0.5 | |
| | Intermediate | arterioles | 7.8 ± 1.8 | 2.0 |
| | | venules | 3.9 ± 0.7 | |
| | Deep | arterioles | 4.1 ± 1.2 | 1.3 |
| | | venules | 3.1 ± 0.7 | |
| ***Monkey L*** | Superficial | arterioles | 9.7 ± 2.1 | 3.7 |
| | | venules | 2.6 ± 1.0 | |
| | Intermediate | arterioles | 6.3 ± 2.5 | 2.7 |
| | | venules | 2.3 ± 0.8 | |
| | Deep | arterioles | 8.3 ± 2.5 | 2.7 |
| | | venules | 3.1 ± 1.8 | |

### 4. Vessel counting

| | *Stimulus Size ($deg^2$)* | *Mean + std (N vessels)* |
|---|---|---|
| ***Monkey E*** | **0.32** | 6.8 ± 7.6 |
| | **0.83** | 16.9 ± 4.4 |
| | **3.39** | 24.7 ± 3.2 |
| | **6.79** | 32 ± 3.3 |
| | **10.18** | 42.5 ± 4.2 |
| | **13.57** | 49.5 ± 3.6 |
| ***Monkey L*** | **0.32** | 13.4 ± 13.2 |
| | **0.83** | 27.8 ± 17.8 |
| | **3.39** | 43.9 ± 19.9 |
| | **6.79** | 52.6 ± 17.8 |
| | **10.18** | 63.7 ± 16.9 |
| | **13.57** | 74.2 ± 14.4 |
